# Regulation of Iron Homeostasis Through Parkin-mediated Lactoferrin Ubiquitylation

**DOI:** 10.1101/2020.06.04.133637

**Authors:** Ankur A. Gholkar, Stefan Schmollinger, Erick F. Velasquez, Yu-Chen Lo, Whitaker Cohn, Joseph Capri, Harish Dharmarajan, William J. Deardorff, Lucy W. Gao, Mai Abdusamad, Julian P. Whitelegge, Jorge Z. Torres

**Author notes:** Corresponding author: Jorge Z. Torres, 607 Charles E. Young Drive East, Los Angeles, CA 90095.

## Abstract

Somatic mutations that perturb Parkin ubiquitin ligase activity and the misregulation of iron homeostasis have both been linked to Parkinson’s disease. Lactotransferrin is a member of the transferrin iron binding proteins that regulate iron homeostasis and increased levels of Lactotransferrin and its receptor have been observed in neurodegenerative disorders like Parkinson’s disease. Here, we report that Parkin binds to Lactotransferrin and ubiquitylates it to regulate iron homeostasis.

---

Parkinson’s Disease (PD) is a debilitating neurodegenerative disease whose incidence has increased over the last decade and it presents a major public healthcare epidemic^1^. The *PARK2* gene that encodes the Parkin E3 ubiquitin ligase is found mutated in familial forms of PD^2^. To better understand the role of Parkin dysfunction in PD, we sought to identify novel Parkin ubiquitylation substrates. First, we established a HEK293 doxycycline-inducible localization and affinity purification (LAP= EGFP-TEV-S-Peptide)-tagged Parkin stable cell line and utilized it to express and tandem affinity purify LAP-Parkin^3^. Eluates were analyzed by mass spectrometry to identify Parkin associating proteins (**Supplementary Fig. 1a**). This analysis identified Parkin (270 peptides) and Lactotransferrin (LTF, 72 peptides) as the most abundant proteins along with tubulin isoforms, proteasome subunits, and CCT/TRiC (Chaperonin containing T-complex/TCP-1 ring complex) subunits (**Supplementary Fig. 1b** and **Supplementary Table 1**).

LTF is a member of the transferrin iron binding proteins that transport iron and regulate intracellular iron levels^4,5^. Increased levels of LTF and its receptor have been reported within nigral neurons in PD patients and in other neurodegenerative disorders^4,6^. Iron homeostasis is important for maintaining normal physiology of neuronal cell populations and iron accumulation leads to neurotoxicity^7^. Due to the importance of LTF in iron homeostasis and its misregulation in PD^4^, we sought to further validate the Parkin-LTF interaction. Reciprocal co-IP experiments from SH-SY5Y neuronal cells and HeLa cells with anti-Parkin and anti-LTF antibodies showed that LTF co-IPd with Parkin and Parkin co-IPd with LTF (**Supplementary Fig. 1c,d,e,f**). Similarly, *in vitro* protein binding reactions with GST-Parkin and FLAG-LTF showed that LTF co-IPd with Parkin (**Supplementary Fig. 1g)**. Together these data indicated that LTF was associating with Parkin.

To understand the significance of the Parkin-LTF association, we asked if LTF was ubiquitylated and whether its ubiquitylation was Parkin-dependent. LAP-LTF was IPd from control siRNA (siCont) or Parkin siRNA (siParkin) treated cells and its ubiquitylation was monitored by immunoblot analysis with anti-ubiquitin antibodies. LAP-LTF was ubiquitylated in the siCont sample and this ubiquitylation was significantly decreased upon Parkin depletion with siParkin (**Fig. 1a**). Next, we asked if LTF was a Parkin substrate using an *in vitro* reconstituted ubiquitylation assay^8^. GST-LTF, GST-Tubulin (positive control) or GST-GFP (negative control) were incubated with an ATP-regeneration system, ubiquitin, E1 ubiquitin-activating enzyme, E2 ubiquitin conjugating enzyme, and WT or LAP-Parkin over expressing HEK293 cell extracts. Parkin substrate ubiquitylation was then monitored by immunoprecipitating the GST-tagged proteins and performing an immunoblot analysis with anti-ubiquitin and anti-GST antibodies. We observed ubiquitylation of GST-LTF and GST-Tubulin (a known substrate of Parkin^9^) as a ladder of increasing molecular weight bands (**Fig. 1b**). Next, we analyzed LTF ubiquitylation reactions by mass spectrometry and determined that LTF was ubiquitylated at 7 different lysine (K) residues with K182 and K649 being the most frequently modified sites (**Supplementary Table 2**). Mapping of the ubiquitylation sites onto the human LTF crystal structure (PDB 1FCK) showed that all sites were on exposed loops (**Fig. 1c**). Next, we analyzed the contribution of the most frequently modified lysines (K182 or K649) to the overall ubiquitylation of LTF by mutating them to alanines and assessing LTF ubiquitylation. LAP-LTF-WT or LTF single (K182A or K649A) or double (K182A/K649A) ubiquitylation site mutants were expressed in HeLa cells, immunoprecipitated, and their ubiquitylation status was monitored using anti-K48 and anti-K63 ubiquitin linkage specific antibodies. Single K182A or K649A mutants showed a reduction in K63-linked ubiquitylation of LTF and ubiquitylation of the LTF K182A/K649A double mutant was highly impaired compared to the WT control (**Fig. 1d**). Together these data indicated that LTF was a Parkin substrate and that K182A and K649A were the most frequently ubiquitylated sites and accounted for the majority of LTF ubiquitylated species.

**Figure 1.**
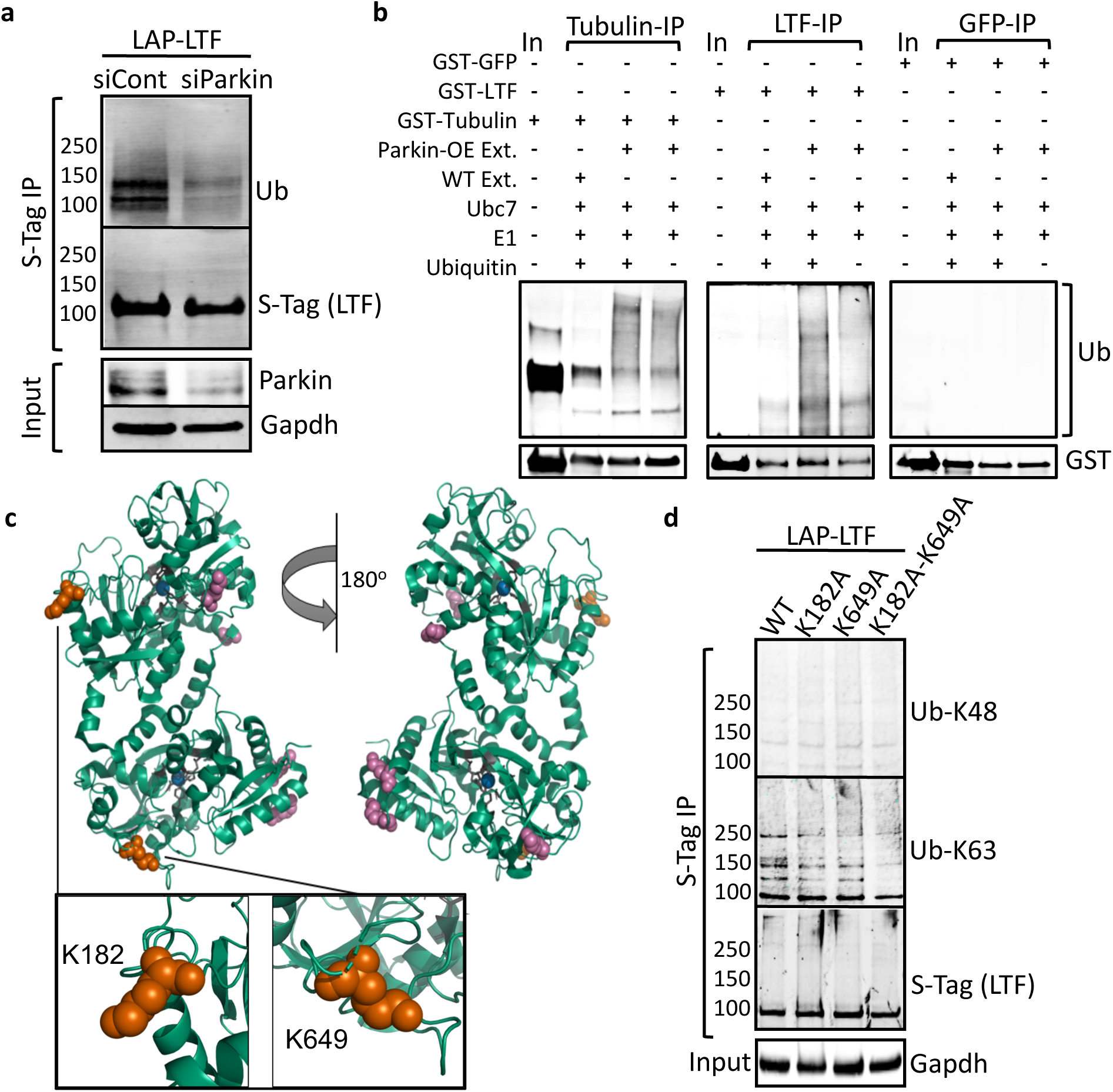
LTF is a substrate of Parkin. (**a**) Immunoblot analysis of LAP-LTF immunoprecipitated from HeLa cells treated with control (siCont) or Parkin (siParkin) siRNA. LAP-LTF ubiquitylation was monitored with anti-ubiquitin antibodies (Ub). Depletion of Parkin was monitored with anti-Parkin antibodies. (**b**) *In vitro* ubiquitylation assays with or without recombinant GST-LTF, GST-Tubulin (positive control) or GST-GFP (negative control); an ATP-regeneration system; ubiquitin; E1 ubiquitin-activating enzyme; E2 ubiquitin conjugating enzyme (Ubc7) and WT or LAP-Parkin over expressing (OE) HEK293 cell extracts (Ext). GST-tagged proteins were immunoprecipitated and their ubiquitylation was monitored with anti-ubiquitin antibodies (Ub). (**c**) LTF ubiquitylation reactions were analyzed by mass spectrometry and the most abundant Parkin-mediated LTF ubiquitylation sites, K182 and K649, were mapped onto the human LTF structure PDB 1FCK (represented as orange spheres). Five additional ubiquitylation sites are represented in magenta spheres. Fe bound to LTF is represented in blue spheres. For a complete list of identified LTF ubiquitylation sites see Supplementary Table 2. (**d**) LAP-LTF-WT or LTF single (K182A or K649A) or double (K182A/K649A) ubiquitylation site mutants were expressed in HeLa cells, immunoprecipitated and their ubiquitylation was monitored using anti-K48 and anti-K63 ubiquitin linkage specific antibodies.!

Next, we sought to determine if ubiquitylation at K182A or K649A was influencing the ability of LTF to regulate intracellular iron levels. Extracts from control HeLa cells or HeLa cells over expressing LTF wild type, K182A single mutant, K649A single mutant, or K182A/K649A double mutant were analyzed for intracellular sulfur, iron and zinc levels using Inductively Coupled Plasma Mass Spectrometry (ICP-MS/MS). Overexpression of the K649A variant alone or the K182A/K649A double mutant resulted in similar, significantly increased intracellular Fe compared to controls, while total sulfur levels and intracellular zinc levels were not altered (**Fig. 2a,b)**. We hypothesized that if Parkin was ubiquitylating LTF on K649A to regulate iron levels, that modulation of Parkin levels would also affect intracellular iron levels. To test this, we performed RNAi-mediated depletion of Parkin levels and again analyzed the extracts for total S, Fe and Zn. Decreasing Parkin levels led to a significant increase, specifically, in Fe content, compared to the controls (**Fig. 2c,d)**. In contrast, extracts from cells that were overexpressing Parkin showed a significant decrease in total Fe, compared to the controls (**Fig. 2e,f)**. Together these data demonstrated that Parkin abundance directly or the mutation of the Parkin-dependent LTF ubiquitylation site (K649A) affect intracellular iron levels, consistent with a role of Parkin regulating iron levels through LTF ubiquitylation.

**Figure 2.**
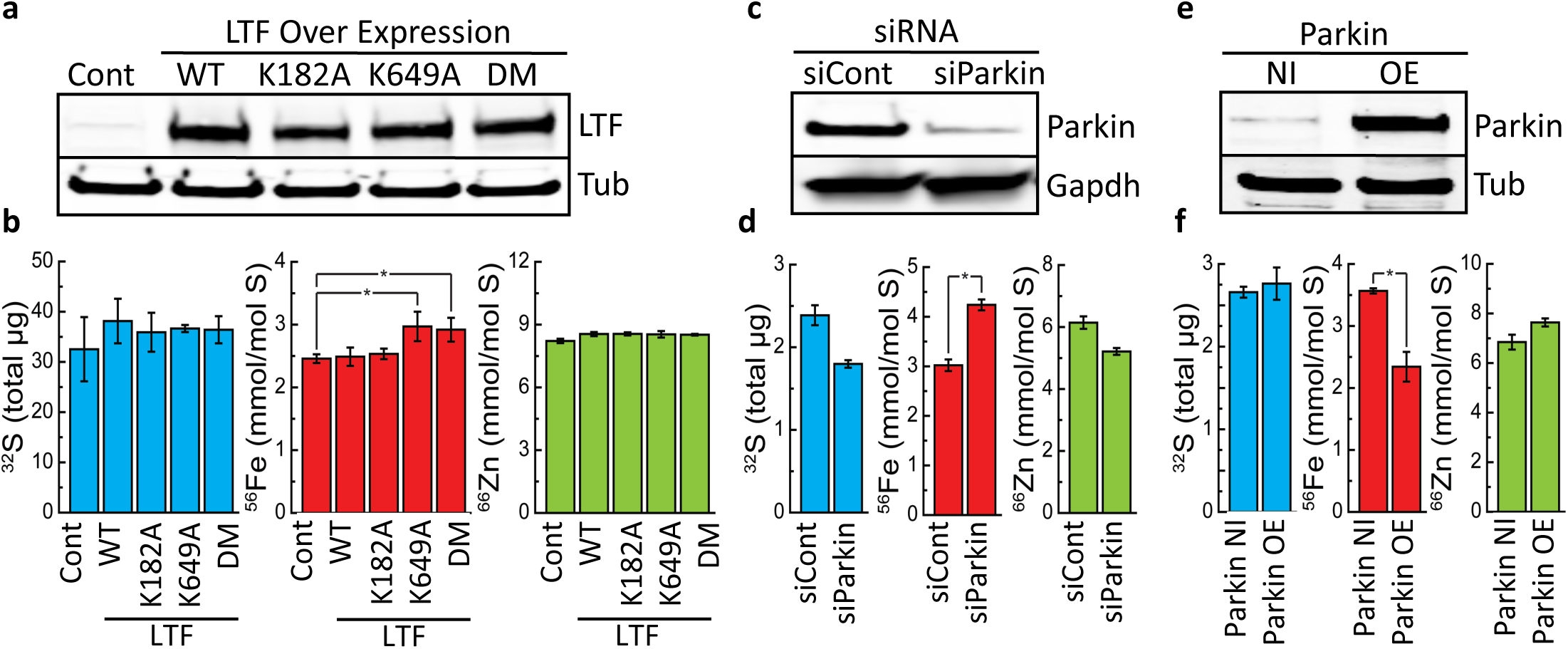
Parkin-mediated ubiquitylation of LTF regulates intracellular iron levels. (**a,b**) Extracts from control HeLa cells (Cont) or HeLa cells over expressing LTF wild type (WT), lysine 182 to alanine (K182A) mutant, lysine 649 to alanine (K649A) mutant, or the double mutant (K182A/K649A) were analyzed for sulfur (^32^S), iron (^56^Fe), and zinc (^66^Zn). (**c,d**) Extracts from HeLa cells transfected with control siRNA (siCont) or Parkin siRNA (siParkin) were analyzed for total sulfur (^32^S), iron (^56^Fe), and zinc (^66^Zn). (**e,f**) Extracts from the HEK293 LAP-Parkin inducible cell line that was not induced (Parkin NI) or induced to over express LAP-Parkin (Parkin OE) were analyzed for total sulfur (^32^S), iron (^56^Fe), and zinc (^66^Zn). (**b,d,f**) Bar graphs show mean ± standard deviation from 3 replicate samples. A *t*-test was used to calculate p values (α < 5%) in the indicated comparisons.

We have discovered a previously undescribed link between Parkin and LTF that influences iron homeostasis. We determined that Parkin binds to and ubiquitylates LTF to regulate intracellular iron levels. Depletion of Parkin led to an increase in intracellular iron levels. Whereas, Parkin overexpression led to a decrease in intracellular iron levels. Importantly, overexpression of a mutant form of LTF, where the ubiquitylation site had been mutated to an alanine (K649A) also led to an increase in intracellular iron levels. We propose that Parkin ubiquitylation of LTF at K649 inhibits LTF’s ability to increase intracellular iron levels and that depletion of Parkin or mutation of K649 on LTF allow LTF to increase intracellular iron levels. The ability of Parkin to regulate iron levels through LTF ubiquitylation may have direct implications to the increased iron levels that are observed in the nigral cells of PD patients.

## METHODS

Methods, including statements of data availability and any associated accession codes and references, are available in the online version of the paper.

## ACKNOWLEDGMENTS

This work was supported by a National Science Foundation Grant NSF-MCB1243645 to J.Z.T., a UCSD/UCLA NIDDK Diabetes Research Center P30 DK063491 grant to J.P.W., a Department of Energy grant DE-FD02-04ER15529 to support S.S., a UCLA Molecular Biology Institute Whitcome Fellowship to E.F.V., and a National Institutes of Health grant GM42143 for instrument support.

## AUTHOR CONTRIBUTIONS

A.A.G. and J.Z.T. contributed to the design, execution and analysis of experiments. M.A., H.D., and W.J.D., contributed to the experimentation. S.S. performed the ICP-MS/MS analyses. E.F.V. and Y-C.L. performed LTF structure modeling. W.C., J.C., L.W.G., and J.P.W. performed mass spectrometry characterization of LTF ubiquitylation. A.A.G. and J.Z.T. wrote the final version of the manuscript.

## ONLINE METHODS

### Cell culture

SH-SY5Y, HeLa and HEK293 Flp-In T-REx LAP-tagged stable cell lines were grown in F12:DMEM 50:50 medium (GIBCO) with 10% FBS, 2 mM L-glutamine and antibiotics, in 5% CO_2_ at 37° C. Cells were induced to express the indicated LAP-tagged proteins by addition of .1 µg/ml doxycycline (Sigma-Aldrich) for the indicated times.

### Plasmids, mutagenesis, and generation of stable cell lines

Full length PARKIN or LTF cDNA was fused to the c-terminus of EGFP (pGLAP1 vector^3^ or FLAG (pCS2-FLAG vector) as described previously^10^. The LTF single (K182A and K649A) and double (K182A/K649A) mutants were generated using the QuickChange Lightning Site-Directed Mutagenesis Kit (Agilent Technologies) with primers carrying the desired mutations. All mutagenesis primers were purchased from Fisher Scientific, see **Supplementary Table 3** for primer sequences. The pGLAP1-PARKIN, pGLAP1-LTF-K182A, pGLAP1-LTF-K649A, and pGLAP1-LTF-K182A/K649A vectors were used to generate doxycycline inducible HeLa Flp-In T-REx stable cell lines that express the fusion proteins from a specific single loci within the genome as described previously^3,11^.

### Immunoprecipitation and LAP purification

The indicted cell lines were harvested and cell extracts were prepared in LAP300 lysis buffer (50 mM Hepes pH 7.4, 300 mM KCl, 1 mM EGTA, 1 mM MgCl_2_, 10% glycerol) plus 0.3% NP40, 0.5 mM DTT, 10 μM MG132, 20 mM NEM and protease and phosphatase inhibitor cocktail (Thermo Scientific). Immunoprecipitations were performed as previously described previously^3^. Samples were resolved in a 4-20% gradient Tris gel (Bio-Rad) with TGS running buffer, transferred to PVDF membrane and immunoblotted with indicated antibodies. Similarly LAP-tag tandem affinity purifications were performed as described previously^3^.

### Identification of Parkin associated proteins by LC-MS/MS

LAP-Parkin KEK293 cells were grown in roller bottles, induced with .1μg/ml Dox, harvested and lysed in the presence of protease (Roche), phosphatase (Pierce), and proteasome inhibitors (MG132, Enzo life sciences). LAP-Parkin was purified from extracts using a tandem affinity purification protocol^3^. Eluates were resolved on a 4-12% SDS-PAG and ten gel slices corresponding to the resolved Parkin purification (and an adjacent control region) were excised and collected, washed twice with 50% acetonitrile, flash frozen, and stored until examination by mass spectrometry. Mass spectrometry-based proteomic analysis was performed at the Harvard Mass Spectrometry and Proteomics Resource Laboratory by microcapillary reverse-phase HPLC nano-electrospray tandem mass spectrometry (µLC/MS/MS) on a Thermo LTQ-Orbitrap mass spectrometer as described previously^12^. The major proteins identified are listed in **Supplementary Table 1**.

### Quantitative metal and sulfur content analysis via ICP-MS/MS

The elemental content was analyzed as described previously^13^, with minor modifications to accommodate the sample material. Briefly, HeLa cells (1×10^8^) were collected by centrifugation at 1,500 *rpm* for 10 minutes. The cells were washed three times in 1 mM Na_2_-EDTA (to remove cell surface– associated metals and remnants from the growth media) and briefly once in Milli-Q water. The cell pellet, after removal of water, was overlaid with 286 µl 70% nitric acid (Fisher, Optima grade, A467-500, Lot 1216040) and digested at room temperature for 24 hours and at 65 °C for 4 hours, before being diluted to a final nitric acid concentration of 2% (v/v) with Milli-Q water. Iron, zinc and sulfur contents of the cell pellets were determined by inductively coupled plasma mass spectrometry (ICP-MS/MS) on an Agilent 8800 Triple Quadrupole ICP-MS instrument, in comparison to an environmental calibration standard (Agilent 5182-4688) and a sulfur standard (Inorganic Ventures CGS1), using ^89^Y as an internal standard (Inorganic Ventures MSY-100PPM). The levels of all analytes were determined in MS/MS mode by quantifying the most abundant isotope (^32^S, ^56^Fe and ^66^Zn); while ^66^Zn was measured directly using He in the collision/reaction cell, ^56^Fe was directly determined using H_2_ as a cell gas and ^32^S was determined via mass-shift from 32 to 48, utilizing O_2_ as a cell gas. The average of 4 technical replicate measurements was used for each individual sample or standard, the average variation in between the technical replicate measurements was 1.1% for all analytes and never exceeded 5% for any individual sample. Triplicate biological replicates were used to determine the variation in between samples, average and standard deviation between biological replicates are depicted in figures. Sulfur content was used to normalize for varying amount of cell material in between samples, since intracellular sulfur levels were unchanged in between the different samples.

### Ubiquitylation reactions

Ubiquitylation reactions were carried out as described previously^14^. Briefly, GST-tagged LTF or positive control GST-tagged Tubulin or negative control GST-tagged-GFP were incubated with or without HEK293 LAP-Parkin extracts, along with an ATP regeneration system, Ubiquitin, E1, and E2 in a buffer containing 20 mM HEPES, 5 mM NaCl, 5 mM MgCl2, DTT, MG132, protease and phosphatase inhibitor cocktail and incubated for 90 minutes at 30° C. GST beads were then added for 30 minutes, washed four times with a wash buffer containing 20 mM HEPES, 100 mM NaCl, 5 mM MgCl2, 15 mM imidazole, 0.5% TritonX, BME, DTT, MG132 and a protease and phosphatase inhibitor cocktail. The beads were then boiled in 2X Laemmli sample buffer (Bio-Rad) and loaded onto a 4-20% TGX gel (Bio-Rad) followed by western transfer. The blots were subsequently probed with anti-GST and anti-Ubiquitin antibodies.

### siRNA treatments

Treatment of cells with ThermoFisher control non-targeting siRNA (12935300) and siRNAs previously shown to deplete PARKIN expression^15^ (1299001-HSS107594/HSS107593) was as described previously^16^.

### Molecular modeling of ubiquitylation sites

Human LTF protein structure was downloaded from the protein database (PDB 1FCK) and the ubiquitylated lysine residues were highlighted using pymol.

### Antibodies

See **Supplementary Table 3** for a list of antibodies used for biochemical purifications, immunoprecipitations, and immunoblotting.

