## Supplemental Material for "Regulation of Iron Homeostasis Through Parkin-mediated Lactoferrin Ubiquitylation"

*Corresponding author:

#### Supplementary Figures

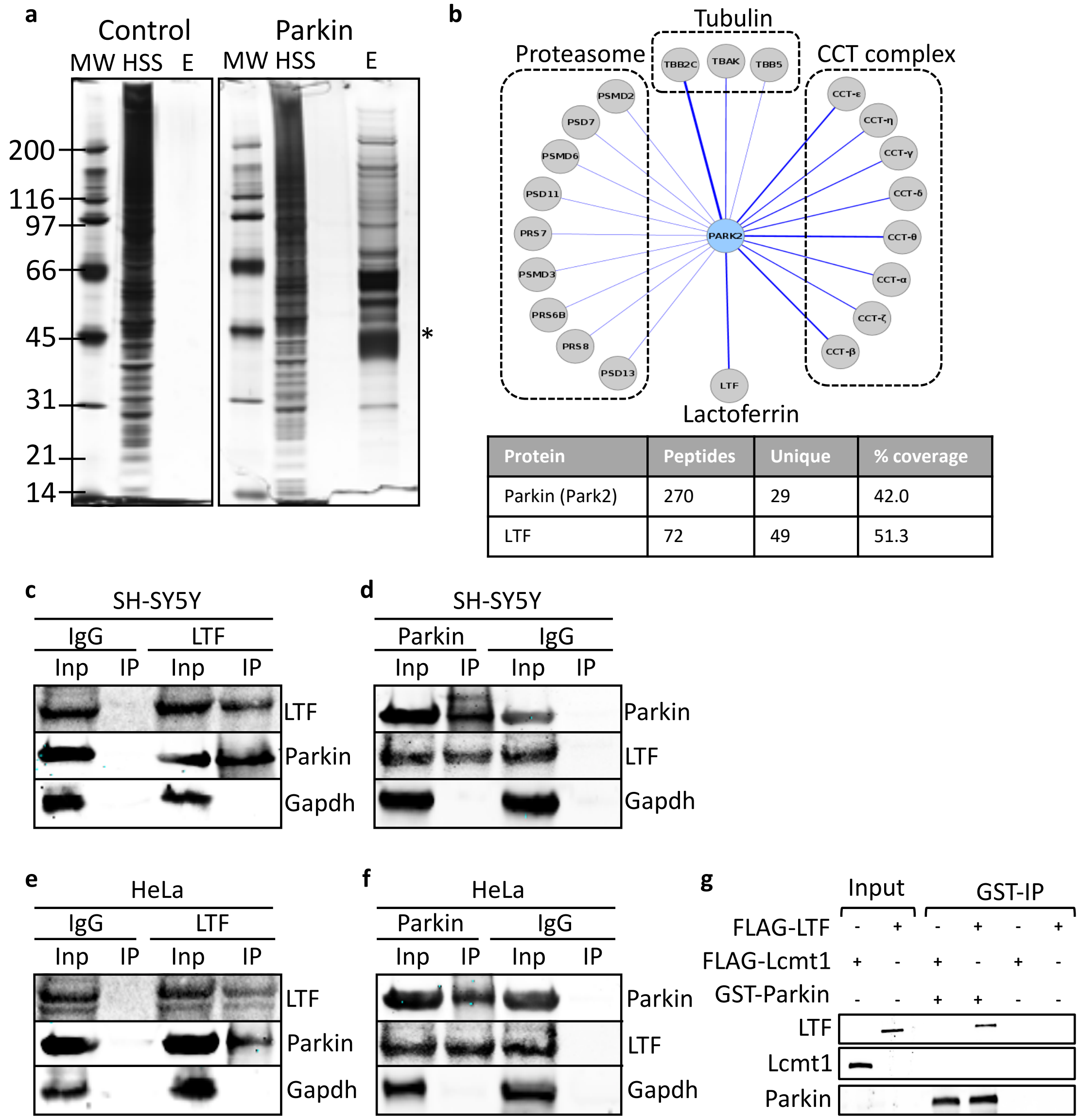


| Protein | Peptides | Unique | % coverage |
|---|---|---|---|
| Parkin (Park2) | 270 | 29 | 42.0 |
| LTF | 72 | 49 | 51.3 |

Supplementary Figure 1. **LTF co-purifies and associates with Parkin.** (**a**) LAP-Parkin tandem affinity purifications. MW= molecular weight marker, HSS= high spin supernatant, E= final eluates. Asterisk denotes the Parkin protein band. (**b**) Cytoscape visualization of Parkin associated proteins, identified by mass spectrometry, showing the major classes of co-purifying proteins. Bottom panel highlights the identification of Lactoferrin (LTF) as a Parkin co-purifying protein. The protein name, number of peptides identified, number of unique peptides, and the percent protein coverage are indicated. See Supplementary Table S1 for a complete list of Parkin co-purifying proteins identified by mass spectrometry. (**c**,**d**,**e**,**f**) SH-SY5Y or HeLa cell extracts were used to perform reciprocal co-immunoprecipitation (Co-IP) experiments using anti-Lactoferrin (LTF), anti-Parkin and control IgG antibodies. Note that endogenous Parkin IPs with endogenous LTF (**c**,**e**) and endogenous LTF IPs with endogenous Parkin (**d**,**f**). (**g**) *In vitro* binding assays performed in the presence or absence of FLAG-LTF, FLAG-Lcmt1, or GST-Parkin. GST-Parkin was IPd and eluates were analyzed by immunoblotting with the indicated antibodies. Note that FLAG-LTF IPs with GST-Parkin, whereas control FLAG-Lcmt1 does not.

**Supplementary Table 1. Summary of LC-MS/MS results from LAP-Parkin purifications**

| Swiss prot ID | Name | # of peptides | Unique | % coverage |
|---|---|---|---|---|
| **Parkin** | | | | |
| O60260 | Park2 | 270 | 27 | 46.9 |
| **Lactoferrin** | | | | |
| P02788 | LTF | 72 | 49 | 51.3 |
| **CCT Complex** | | | | |
| P17987 | CCT-α | 49 | 22 | 39.2 |
| P78371 | CCT-β | 74 | 24 | 39.7 |
| P49368 | CCT-γ | 49 | 24 | 35.8 |
| P50991 | CCT-δ | 48 | 17 | 32.3 |
| P02788 | CCT-ε | 71 | 25 | 35.1 |
| P40227 | CCT-ζ | 46 | 19 | 28.5 |
| Q99832 | CCT-η | 46 | 21 | 33.5 |
| P50990 | CCT-θ | 82 | 29 | 48.6 |
| **Proteasome Complex** | | | | |
| Q15008 | PSMD6 | 6 | 6 | 17.2 |
| P62195 | PRS8 | 3 | 3 | 10.1 |
| O00231 | PSD11 | 2 | 2 | 5.5 |
| P43686 | PRS6B | 2 | 2 | 3.8 |
| P35998 | PRS7 | 3 | 2 | 6 |
| Q9UNM6 | PSD13 | 2 | 2 | 5.3 |
| O43242 | PSMD3 | 3 | 3 | 5.8 |
| Q13200 | PSMD2 | 2 | 2 | 3 |
| p51665 | PSD7 | 1 | 1 | 3.7 |
| **Tubulins** | | | | |
| P68363 | TBAK | 41 | 12 | 25.7 |
| P68371 | TBB2C | 111 | 17 | 38.4 |
| P07437 | TBB5 | 10 | 2 | 6.1 |
| **Ubiquitin** | | | | |
| P62988 | Ubiquitin | 22 | 4 | 61.8 |
| **Chaperones** | | | | |
| P07900 | HSP90a | 22 | 12 | 17.4 |
| P08238 | HSP90b | 4 | 3 | 4.3 |
| P11142 | HSP70/HSP7[illegible] | 130 | 29 | 40.2 |
| P08107 | HSP70.1 | 3 | 1 | 1.7 |
| Q27965 | HSP70.2 | 73 | 19 | 27.3 |
| P11021 | HSP70/GRP7[illegible] | 16 | 12 | 22.6 |
| P38646 | HSP70/GRP7[illegible] | 11 | 7 | 12.5 |
| P10809 | HSP60 | 21 | 14 | 20.2 |
| O95816 | BAG2 | 9 | 6 | 25.6 |
| P31689 | DNJA1 | 12 | 7 | 19.1 |
| O60884 | DNJA2 | 12 | 7 | 16 |
| Q9Y2Z0 | SUGT1 | 3 | 3 | 11 |

### Supplementary Table 2. LTF Ubiquitylated Peptides

**Analysis 1**

| Query | Start | End | Observed | Mr(expt) | Mr(calc) | ppm | M | Score | Expect | Rank | Peptide | Peptides | LTF isoform 1 NP_002334.2 |
|---|---|---|---|---|---|---|---|---|---|---|---|---|---|
| 11691 | 119 | 139 | 597.5474 | 2386.1603 | 2386.2175 | -24 | 2 | 6 | 6.4 | 10 | K.KGGSFQLNELQGLKSCHTGLR.R + GlyGly (K) | 1 of 11 | K119 |
| 11513 | 171 | 190 | 753.353 | 2257.0371 | 2257.0408 | -1.67 | 1 | 7 | 0.77 | 1 | R.FFSASCVPGADKGQFPNLCR.L + GlyGly (K) | 2 of 2 | **K182** |
| 11514 | 171 | 190 | 753.3554 | 2257.0444 | 2257.0408 | 1.57 | 1 | 11 | 0.5 | 1 | R.FFSASCVPGADKGQFPNLCR.L + GlyGly (K) | 2 of 2 | **K182** |
| 3340 | 292 | 299 | 525.2677 | 1048.5208 | 1048.5301 | -8.86 | 1 | 3 | 2.9 | 7 | R.QAQEKFGK.D + GlyGly (K) | 1 of 1 | K296 |
| 10237 | 300 | 315 | 932.989 | 1863.9635 | 1863.9479 | 8.35 | 2 | 8 | 2.6 | 7 | K.DKSPKFQLFGSPSGQK.D + GlyGly (K) | 1 of 1 | K305 |
| 11081 | 642 | 658 | 678.2952 | 2031.8637 | 2031.8666 | -1.46 | 1 | 11 | 0.18 | 1 | R.NGSDCPDKFCLFQSETK.N + GlyGly (K) | 1 of 1 | **K649** |

**Analysis 2**

| Query | Start | End | Observed | Mr(expt) | Mr(calc) | ppm | M | Score | Expect | Rank | Peptide | Peptides | LTF isoform 1 NP_002334.2 |
|---|---|---|---|---|---|---|---|---|---|---|---|---|---|
| 9491 | 171 | 190 | 753.3558 | 2257.0455 | 2257.0408 | 2.06 | 1 | 1 | 1 | 2 | R.FFSASCVPGADKGQFPNLCR.L + GlyGly (K) | 4 of 4 | **K182** |
| 9493 | 171 | 190 | 753.358 | 2257.0523 | 2257.0408 | 5.06 | 1 | 10 | 0.83 | 1 | R.FFSASCVPGADKGQFPNLCR.L + GlyGly (K) | 4 of 4 | **K182** |
| 9494 | 171 | 190 | 753.3588 | 2257.0546 | 2257.0408 | 6.11 | 1 | 13 | 0.54 | 1 | R.FFSASCVPGADKGQFPNLCR.L + GlyGly (K) | 4 of 4 | **K182** |
| 9496 | 171 | 190 | 753.36 | 2257.0581 | 2257.0408 | 7.65 | 1 | 9 | 0.49 | 2 | R.FFSASCVPGADKGQFPNLCR.L + GlyGly (K) | 4 of 4 | **K182** |
| 8077 | 520 | 535 | 603.2715 | 1806.7928 | 1806.7876 | 2.87 | 0 | 16 | 0.54 | 1 | R.SNLCALCIGDEQGENK.C + GlyGly (K) | 1 of 1 | K535 |
| 9059 | 642 | 658 | 1016.944 | 2031.8735 | 2031.8666 | 3.39 | 1 | 18 | 0.061 | 1 | R.NGSDCPDKFCLFQSETK.N + GlyGly (K) | 3 of 3 | **K649** |
| 9060 | 642 | 658 | 678.2996 | 2031.8769 | 2031.8666 | 5.04 | 1 | 9 | 0.13 | 1 | R.NGSDCPDKFCLFQSETK.N + GlyGly (K) | 3 of 3 | **K649** |
| 9061 | 642 | 658 | 678.2997 | 2031.8774 | 2031.8666 | 5.31 | 1 | 15 | 0.084 | 1 | R.NGSDCPDKFCLFQSETK.N + GlyGly (K) | 3 of 3 | **K649** |
| 8110 | 695 | 709 | 905.9499 | 1809.8852 | 1809.8753 | 5.48 | 1 | 4 | 2.7 | 4 | K.KCSTSPLLEACEFLR.- + GlyGly (K) | 1 of 3 | K695 |

**Combined**

| Peptide | Peptides | LTF isoform 1 |
|---|---|---|
| K.KGGSFQLNELQGLKSCHTGLR.R + GlyGly (K) | 1 of 11 | K119 |
| R.FFSASCVPGADKGQFPNLCR.L + GlyGly (K) | 6 of 6 | **K182** |
| R.QAQEKFGK.D + GlyGly (K) | 1 of 1 | K296 |
| K.DKSPKFQLFGSPSGQK.D + GlyGly (K) | 1 of 1 | K305 |
| R.SNLCALCIGDEQGENK.C + GlyGly (K) | 1 of 1 | K535 |
| R.NGSDCPDKFCLFQSETK.N + GlyGly (K) | 4 of 4 | **K649** |
| K.KCSTSPLLEACEFLR.- + GlyGly (K) | 1 of 3 | K695 |

**Supplementary Table 3. Reagents Used**

**1. Primers**

| Name | Forward Primer | Reverse Primer | Company |
|---|---|---|---|
| LTF | 5'GGGGACAAGTTTGTACAAAAAAGCAGGCTTCGAAGGAGATAGAACCATGGGGAAACTTGTCTTCCTCGTCCTGC3' | 5'GGGGACCACTTTGTACAAGAAAGCTGGGTCTCACTTCCTGAGGAATTCACAGGC3' | ThermoFisher |
| LTF (K182A) | 5'CTGTGTTCCCGGTGCAGATGCAGGACAGTTCCCCAA3' | 5'TTGGGGAACTGTCCTGCATCTGCACCGGGAACACAG3' | ThermoFisher |
| LTF (K649A) | 5'GACTGGAATAAGCAAAACGCGTCCGGGCAGTCAGATCC3' | 5'GGATCTGACTGCCCGGACGCGTTTTGCTTATTCCAGTC3' | ThermoFisher |
| LTF (K182A-K649A) | Used K649A primers on LTF (K182A) mutant | Used K649A primers on LTF (K182A) mutant | ThermoFisher |
| Parkin | 5'GGGGACAAGTTTGTACAAAAAAGCAGGCTTCGAAGGAGATAGAACCATGGGGATAGTGTTTGTCAGGTTC3' | 5'GGGGACCACTTTGTACAAGAAAGCTGGGTCCTACACGTCGAACCAGTG3' | ThermoFisher |
| LCMT-1 | 5'GGGGACAAGTTTGTACAAAAAAGCAGGCTTCGAAGGAGATAGAACCATGGGGCTTCCCTGTGCAAGAGAACTCC3' | 5'GGGGACCACTTTGTACAAGAAAGCTGGGTCCTAATAAGTTATCTCCTTCAGC3' | ThermoFisher |

**2. siRNA**

| Taget | siRNA ID | Catalog # | Company |
|---|---|---|---|
| Negative Control | N/A | 12935300 | ThermoFisher |
| PARK2 | HSS107594 | 1299001 | ThermoFisher |
| PARK2 | HSS107593 | 1299001 | ThermoFisher |

**3. Antibodies**

| Target | Name | Catalog # | Company |
|---|---|---|---|
| Ubquitin-K63 | Ubquitin (linkage-specific K63) [EPR8590-448] | Ab179434 | Abcam |
| Ubquitin-K48 | Ubquitin (linkage-specific K48) [EP8589] | Ab140601 | Abcam |
| LTF | Lactoferrin [2B8] | Ab10110 | Abcam |
| Gapdh | Gapdh | GTX100118 | GeneTex |
| Ubquitin | Mono- and polyubiquitinylated conjugates monoclonal antibody (FK2) | BML-PW8810-0100 | Enzo |
| Parkin | Parkin | Ab15954 | Abcam |
| Parkin | Parkin | #2132 | Cell Signaling |
| S-Tag | S tag | GTX19321 | GeneTex |
| GST | GST Tag (3G12B10) | 66001-2-Ig | Proteintech |
| IgG | normal rabbit IgG | 12-370 | Sigma Aldrich |
| IgG | normal mouse IgG | sc-2025 | Santa Cruz |

**4. Proteins**

| Name | Details | Catalog # | Company |
|---|---|---|---|
| LTF | GST-Lactoferrin, Human | H00004057-P01-2ug | Novus |
| Tubulin | GST-Tubulin, Human | SRP5148-20UG | Sigma |
| Ubc7 | His-UBE2G1 (UBC7), Human | 009-001-U35S | Rockland |
| E1 | His-E1, Human | BML-UW9410-0050 | Enzo |
| Ubquitin | His-Ubiquitin, Human | BML-UW8610-0001 | Enzo |
| GFP | GST-GFP | Torres Lab | Torres Lab |

**5. Affinity Resins**

| Name | Details | Catalog # | Company |
|---|---|---|---|
| Protein A Beads | Affi-Prep Protein A Media | 156-0005 | BIO-RAD |
| Protein G Beads | Pierce™ Protein G Magnetic Beads | 88847 | ThermoFisher |
| GST Beads | Pierce™ Glutathione Magnetic Agarose Beads | 78601 | ThermoFisher |
| S Beads | S-protein Agarose | 69704-4 | Millipore |
